# Development of plant regeneration and *Agrobacterium tumefaciens*-mediated transformation methodology for *Physalis pruinosa*

**DOI:** 10.1101/386235

**Authors:** Kerry Swartwood, Joyce Van Eck

## Abstract

*Physalis pruinosa*, also known as groundcherry, produces a small, yellow, highly nutritious edible fruit that is enveloped by a papery husk. In order for the potential of large-scale production of *P. pruinosa* fruit to be realized, undesirable characteristics, such as an unmanageable, sprawling growth habit and extensive fruit drop, need to be improved by exploiting approaches available through plant breeding, genetic engineering, and gene editing. In this study, we established plant regeneration and *Agrobacterium tumefaciens*-mediated methods to allow application of genetic engineering and gene editing of *P. pruinosa*. Cotyledon and hypocotyl explants from 7 – 8-day-old *in vitro*-grown seedlings were assessed for plant regeneration. Explants were cultured for 2 weeks on a Murashige and Skoog salts-based medium that contained 2 mg/L zeatin followed by transfer to medium containing 1 mg/L zeatin. Only hypocotyl explants regenerated shoots. Hypocotyl explants were infected with *Agrobacterium tumefaciens* strain AGL1 containing the pJL33 binary vector that has the green fluorescent protein (*GFP*) reporter and neomycin phosphotransferase II (*nptII*) selectable marker genes. After cocultivation, explants were cultured on selective plant regeneration medium that contained 50, 100, 200, 250, and 300 mg/L kanamycin to determine the most effective level for efficient recovery of transgenic lines. Based on rooting of regenerated shoots on selective medium, GFP visualization, and PCR analysis for the presence of the *nptII* gene, medium containing 200 mg/L kanamycin resulted in the highest transformation efficiency at 24%. This study sets the foundation for future genetic engineering and gene editing approaches for improvement of *P. pruinosa*.

## Introduction

The *Solanaceae* family is comprised of a very diverse group of approximately 3,000 species that represent important food crops, ornamentals, and plants with medicinal properties. There is potential for some *Solanaceae*, including those in the *Physalis* genera, as new specialty fruit crops, especially in temperate regions, however, trait modification for desirable agronomic characteristic would be needed for agricultural production, (Valdivia-Mares et al. 2016). Within the *Solanaceae*, *Physalis* is one of five genera that displays a unique morphological attribute in the form of a papery husk, also referred to as an inflated calyx, that results when sepals elongate and completely envelop the developing fruit (Wang et al. 2015). The husk provides protection from diseases and pests and has also been shown to play a role in carbohydrate translocation and fruit development (Fischer and Ludders 1997).

*Physalis* most recognizable as food crops are groundcherry (*P. pruinosa*), goldenberry (*P. peruviana*), and tomatillo (*P. ixocarpa*). *P. pruinosa* originated in Eastern North America, whereas *P. peruviana* and *P. ixocarpa* originated in Peru and Mexico, respectively (Singh et al. 2014). All three of these *Physalis* species are primarily consumed as fresh fruit, but *P. pruinosa* and *P. peruviana* are also processed to make jams, juices, raisins, and snack products (Puente et al. 2011).

*P. pruinosa* is of special interest for our work because of its potential as a specialty fruit crop that could provide farmers, namely in temperate regions based on its origins, with an additional source of income that would be further spurred by the growing consumer demand for new and unique fruits (Singh et al. 2014). One advantage of *P. pruinosa* from an agricultural standpoint is it can be grown in a wide variety of soil types with successful crop production even in poor, sandy conditions (Wolff 1991). However, wider adoption as a food crop would be more likely realized through improvements of undesirable agronomic characteristics that would negatively impact agricultural production, especially on a large scale. One such undesirable characteristic, is its sprawling growth habit that would make it unmanageable in production. Another trait for improvement would be fruit size because *P. pruinosa* fruit are very small at approximately 1 g and 1 cm in diameter. The fruit ranges from greenish-yellow to golden when ripe depending on the variety and it has a sweeter flavor than tomato, which is another member of the *Solanaceae*. The fruit are encased in a husk or inflated calyx that forms from elongation of the sepals and turns papery when the fruit are ripe. *P. pruinosa* is often referred to as groundcherry because the fruit drops to the ground as a result of an abscission zone, similar to the joint found in some tomato pedicels, making harvest difficult (Lee et al. 2018). In addition to the negative aspect of harvestability, gathering fruit from the ground also imposes a food safety risk that could lead to food-borne illness.

Improvement of the less desirable characteristics of *P. pruinosa*, such as growth habit, small fruit size, and fruit drop, would help to increase the likelihood of its broader adoption as a specialty fruit crop. Unfortunately, there is a lack of gene function and genomic information for *P. pruinosa* that are necessary to develop improvement programs. Expansion of resources is critical for further development of this underutilized solanaceous species as a viable food crop. Perhaps knowledge gained from our work on gene identification, function and trait modification could be applied to improve agronomic characteristics of other underutilized or orphan crops to enhance their agricultural productivity and contribute towards diversification of our food supply especially in regions where people rely on a limited number of sources. Therefore, our efforts have been focused on generation of resources, including a gene delivery method reported here, that will allow these improvements to be put into practice (Lemmon et al. 2018). Our intent for this study was to develop *Agrobacterium tumefaciens*-mediated transformation methodology for *P. pruinosa* to facilitate approaches for improvement that include functional studies and application of gene editing technology. In brief, we found that infection of hypocotyl explants from *in vitro*-grown seedlings was a viable approach for recovery of transgenic lines. To our knowledge, there are no reports on transformation methods for *P. pruinosa*.

## Materials and Methods

### Plant material and in vitro seed germination

Seeds of *Physalis pruinosa* were obtained from the Solanaceae Germplasm Bank at the Botanical Garden of Nijmegen. To increase the number of seeds to provide the necessary amount for development of transformation methods, seeds were directly sown and germinated in soil in 72-cell inserts in plastic flats that were placed on heat mats in the greenhouse and grown under long-day conditions (16-h light/8-h dark) under natural light supplemented with artificial light from high-pressure sodium bulbs (~250 µmol m^-2^ s^-1^). Daytime and nighttime temperatures were 26–28 **°**C and 18–20 **°**C, respectively, with a relative humidity of 40-60%. Mature fruits were collected from greenhouse-grown *P. pruinosa* plants. To remove the mucilage gel surrounding the seeds extracted from fruit, seeds were pressed against a metal strainer, placed into mesh extraction bags, and soaked in Rapidase C80 KPO (Centerchem.com) for 1 - 1.5 hours. The bag was thoroughly rinsed with water before soaking in 40% bleach for 10 min followed by 5 - 7 rinses, and air-dried at room temperature (approximately 72°C) on paper towels.

Seeds for regeneration and transformation experiments were surface sterilized in 20% (v/v) bleach solution containing Tween-20 for 20 min with gentle agitation followed by 3 rinses in sterile deionized water. Surface sterilized seeds were germinated in Magenta™ GA-7 boxes (from here forward referred to as Magenta boxes) containing 50 mL of germination medium composed of 2.15 g/L Murashige and Skoog (MS) (Murashige and Skoog 1962) salts (Caisson Labs), 100 mg/L myo-inositol, 2 mg/L thiamine, 0.5 mg/L pyridoxine, 0.5 mg/L nicotinic acid, 10 g/L sucrose and 8 g/L agar (Sigma-Aldrich, St. Louis, MO). The pH of the medium was adjusted to 5.8.

### Plant regeneration assessment

Approximately 7 - 8 days after the seeds were placed on germination medium, before the first true leaves emerged (Fig. 1), cotyledon and hypocotyl explants were excised. To prepare explants, seedlings were placed on a sterile paper towel moistened with sterile deionized water. Cotyledons were excised at the petioles and cut into approximately 0.25 - 0.5 cm cross sections. Hypocotyl explants were prepared by cutting immediately below the shoot apex, removing roots, and cutting the remaining hypocotyl into 0.5 - 1 cm sections.

**Figure 1.**
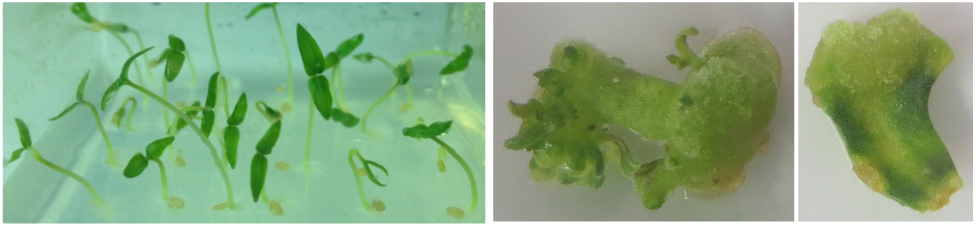
Response of *Physalis pruinosa* hypocotyl and cotyledon explants after 5 weeks on plant regeneration medium (2 wks on 2Z, 3 wks on 1Z). Left: Seven-day-od, *in vitro*-grown seedlings used as a source of cotyledon and hypocotyl explants. Center: Shoot regeneration from hypocotyl explants. Right: Callus formation on cotyledon explants.

Cotyledon and hypocotyl explants were placed onto first phase regeneration medium designated 2Z, which contained 4.3 g/L MS salts, 100 mg/L myo-inositol, 1 mL/L of a 1000X solution of modified Nitsch and Nitsch vitamins (per 100 mL: glycine 0.2 g, nicotinic acid 1 g, pyridoxine-HCl 0.05 g, thiamine-HCl 0.05 g, folic acid 0.05 g, d-biotin 0.004 g, pH 7.0), 20 g/L sucrose, 2 mg/L trans-zeatin, and 5.2 g/L TC Gel (Caisson). Cotyledons were placed adaxial side down on the medium. After 2 weeks, coytledon and hypocotyl explants were transferred to second phase plant regeneration medium, 1Z, which contained the same components as 2Z except that the concentration of trans-zeatin was decreased to 1 mg/L. Petri plates were used for the first transfer to 1Z, but subsequently explants were transferred to fresh media in either petri plates or Magenta boxes depending on the size of the shoots.

When shoots were 2 cm tall they were excised and transferred to rooting medium designated RM, which contained 4.3 g/L MS salts, 1 mL/L of a modified Nitsch and Nitsch vitamins solution (see above), 30 g/L sucrose, 8 g/L Difco Bacto agar (Becton, Dickinson and Company, Franklin Lakes, NJ). Magenta boxes that contained 62.5 mL of RM were used for the rooting phase.

### Agrobacterium strain and binary vector

The pJL33 binary vector (Floss et al. 2013) was introduced by electroporation into electrocompetent *Agrobacterium tumefaciens* AGL1 and referred to as AGL1/JL33 (Lazo et al. 1991). pJL33 contains a synthetic version of the gene for green fluorescent protein designated *sGFP (S65T)* (Chiu et al. 1996) and the neomycin phosphotransferase (*nptII*) selectable marker gene, which confers resistance to kanamycin. The Cauliflower Mosaic Virus 35S promoter (CaMV35S) drives expression of *sGFP (S65T)* and *nptII* in pJL33 (Fig. 2).

**Figure 2.**
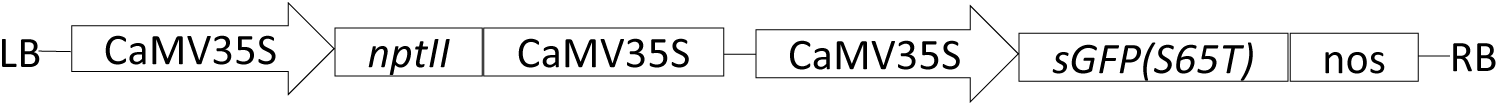
T-DNA region of the binary vector pJL33 used for *Physalis pruinosa* transformation. The T-DNA region contains the antibiotic selection marker *nptII* driven by the CaMV35S promoter. The CaMV35S promoter also drives expression of a synthetic *GFP* gene, *sGFP (S65T)* (Chiu et al. 1996).

To prepare for transformation experiments, AGL1/pJL33 was streaked from a previously prepared glycerol stock maintained at −80°C onto solidified MGL selective medium (Per liter: 5 g tryptone, 2.5 g yeast extract, 5g NaCl, 5 g mannitol, 0.1 g MgSO_4_, 0.25 g K_2_HPO_4_, 1.2 g glutamic acid, 15 g sucrose, 50 mg/L carbenicillin and 50 mg/L kanamycin; pH was adjusted to 7.2, 15 g Bacto Agar) and maintained at 28°C for 48 hours. Four single, well-formed colonies were selected, transferred to 50 mL of liquid LB selective medium, and maintained in a shaking incubator at 28°C for 18 – 24 hrs at 250 rpm or the length of time needed to reach an OD_600_ of 0.6 - 0.7. The *Agrobacterium* suspension was centrifuged at 8000 rpm for 10 min at 20°C. The pellet was resuspended in 50 mL of 2% MSO medium (4.3 g/L MS salts, 100 mg/L myo-inositol, 0.4 mg/L thiamine, and 20 g/L sucrose, pH to 5.8) by vortexing.

### Plant transformation and optimization of kanamycin concentration

One day prior to infection with AGL1/pJL33, hypocotyl explants were excised from 7 - 8-day-old *P. pruinosa* seedlings (before emergence of first true leaves) and cultured on 2Z medium. For infection, the explants were incubated in the *Agrobacterium*/2% MSO suspension for 5 min then transferred to a sterile paper towel to briefly drain excess suspension. The explants were placed back onto 2Z and cultured in the dark at 19°C for 48 hrs. Plates were sealed with parafilm for the cocultivation period.

After 48 hrs, the explants were transferred to selective 2Z (designated 2ZK) that contained 300 mg/L timentin and kanamycin at experimental levels of 50, 100, 200, 250, and 300 mg/L. Two weeks later the explants were transferred to selective 1ZK medium that contained the same concentrations of timentin and kanamycin as 2ZK for testing kanamycin sensitivity. Explants were transferred to freshly prepared 1ZK medium every two weeks in either Petri plates or Magenta boxes depending upon size of shoots regenerating from the explants.

When regenerated shoots were approximately 2 cm tall, they were excised from explants and transferred to selective RM (designated RMK) that contained 100 mg/L kanamycin and 300 mg/L timentin in Magenta boxes. For acclimation to greenhouse conditions, culture medium was washed from well-rooted plants before transfer to soil. The plants were immediately covered with plastic domes, which were gradually removed over the course of 5 days. Plants were maintained in a growth chamber for 2 – 3 weeks then transferred to a greenhouse.

Unless otherwise noted, the following conditions were followed. The pH of all media was adjusted to 6.0 before autoclaving. For all media, trans-zeatin, kanamycin, and timentin were dispensed from filter sterilized stock solutions into autoclaved medium cooled to 55°C. The size of the Petri plates used was 100 mm x 20 mm. Plates were sealed with Micropore tape (Fisher Scientific, Hampton, NH). All cultures were maintained at 24 ± 2°C under a 16 h light/8 h dark photoperiod at 57 – 65 uE m^-2^ s^-1^.

### Green fluorescent protein visualization

*In vitro* material at different time points post infection with AGL1/pJL33 through the stages of young plants, fruit, and seeds, were assessed for green fluorescent protein (GFP) expression. Comparisons of putative transgenic material were made with control material (non-infected). All material was examined with an Olympus SZX-12 stereo microscope equipped for fluorescence imaging. To visualize GFP expression, an LP Green filter cube was used that had an excitation filter of 470/40 and emission filter of LP 500. Images were captured with a color CCD camera connected to the ProgRes C14 acquisition software.

### DNA extraction and PCR analysis

Leaf material was collected from plants that rooted on RMK and exhibited GFP expression for DNA isolation to facilitate PCR analysis to confirm the presence of the *nptII* selectable marker gene. In brief, DNA was extracted according to a standard cetyl-trimethyl-ammonium bromide (CTAB) protocol. Following maceration of the tissue in a CTAB buffer, the suspension was treated with chloroform/isoamyl alcohol (24:1) and DNA was precipitated by the addition of iso-propanol followed by incubation at −80C for 10 min.

Primers used to detect *nptII* were forward 5’-GGC TGG AGA GGC TAT TC-3’ and reverse 5’-GGA GGC GAT AGA AGG CG-3’. The diagnostic amplicon size expected with these primers is approximately 735 bp. The PCR program started with a one-step cycle of 3 min at 94°C, followed by 34 cycles of 30 s at 94°C, 30 s at 58°C, 45 s at 72°C. DNA was separated and visualized by electrophoresis through a 1.5% agarose, ethidium bromide-stained gel.

## Results

### Plant regeneration from cotyledons and hypocotyls

*P. pruinosa* seedlings were at the cotyledon (pre-true leaf) stage 7 - 8 days after seeds were cultured on germination medium (Fig. 1). Approximately 85% of the seeds germinated. To test plant regeneration potential from seedling material, cotyledon and hypocotyl segments were cultured on a first phase plant regeneration medium, 2Z, for 2 wks, then the second phase, 1ZK. Callus developed along the cut edges of the cotyledon and hypocotyl sections, however, shoots only developed on hypocotyl explants (Fig. 1). Shoots were first observed after approximately 3 weeks on 1Z medium.

### Transformation and optimization of kanamycin concentration for selection and rooting of transgenic lines

A set of explants that did not undergo infection was cultured on 2Z followed by 1Z as positive controls. A second set was cultured on 2ZK and 1ZK at each concentration of kanamycin to serve as negative controls. Plant regeneration was observed on all positive controls. As for the negative controls, shoot development decreased as the concentration of kanamycin increased.

Following the 2-day cocultivation of infected material, 25 hypocotyl explants were cultured per Petri plate of 2ZK with 3 replicates per kanamycin concentration (50, 100, 200, 250, and 300 mg/L) to determine the level that would result in the highest transformation efficiency. Subsequent transfers were done to 1ZK containing the corresponding kanamycin concentrations. Experiments were performed 2 separate times. Initiation of shoot regeneration was observed approximately 3 wks after transfer to 1ZK. As the concentration of kanamycin increased from 50 to 300 mg/L kanamycin, the number of regenerated plants decreased (Fig. 3). Shoots were transferred to RMK and roots initiated after 5 – 9 days after transfer. We observed that a large number of shoots recovered from hypocotyl explants on 50 and 100 mg/L did not root as compared to those recovered from culturing on medium that contained 200 – 300 mg/L kanamycin. Based on this result, we concluded that 50 and 100 mg/L of kanamycin did not prevent uninfected cells from developing into plants.

**Figure 3.**
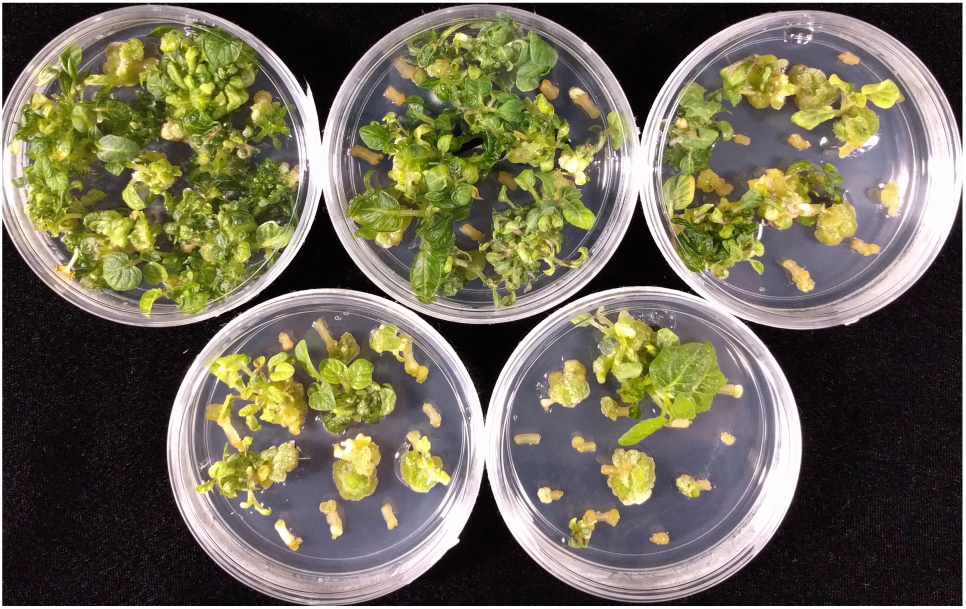
Determination of an effective kanamycin concentration for selection of transgenic lines from *Physalis pruinosa* hypocotyl explants infected with *Agrobacterium* tumefaciens containing pJL33. Top row, kanamycin concentration (mg/L) left to right: 50, 100, 200. Bottom row left to right: 250, 300.

We calculated transformation efficiency for each concentration by dividing the number of plants that rooted on RMK by the total number of hypocotyl explants cultured on each concentration of kanamycin (Table 1). Only 1 shoot was removed per hypocotyl explant to ensure the recovery of independent transgenic events. The highest efficiency resulted from infected material cultured on 2ZK and 1ZK that contained 200 mg/L kanamycin. PCR analysis and assessment for GFP expression was done on all rooted shoots to further verify they were transgenic.

**Table 1.** Effect of kanamycin concentration on transformation efficiency of *Physalis pruinosa*.

| Kanamycin<br>(mg/l) | Total number of<br>hypocotyl explants | Total number of<br>rooted shoots<sup>a</sup> | Average transformation<br>efficiency (±) SE<sup>b</sup> |
| --- | --- | --- | --- |
| 50 | 247 | 9 | 9.4 ± 4.30 |
| 100 | 152 | 14 | 9.3 ± 4.20 |
| 200 | 247 | 33 | 23.6 ± 4.11 |
| 250 | 152 | 22 | 14.4 ± 0.95 |
| 300 | 152 | 21 | 13.7 ± 1.05 |

<sup>a</sup>Shoots rooted on RMK.
<sup>b</sup>Transformation efficiency was calculated as percent of stable transgenic lines recovered from the number of hypocotyl explants infected with *Agrobacterium tumefaciens*. Efficiency values shown are the average from 2 experiments ± the standard error (SE) calculated from 6 biological replicate, with the exception of values for 50 and 200 mg/l, which represent 3 experiments and 9 biological replicates.

### Assessment of transgenesis by GFP visualization and PCR analysis

We chose the binary vector pJL33 (Floss et al. 2013) (Fig. 2) for development of *P. pruinosa* transformation methodology because it contains a reporter gene cassette for the green fluorescent protein (GFP), which provides a non-destructive screening approach that allowed us to monitor the progress of transformation post-infection through mature plants in the greenhouse, fruit, and seeds. pJL33 contains a modified version of the *GFP* gene from the jellyfish (*Aequorea victoria*) that results in a 20-fold higher expression of GFP in plant cells because of codon usage optimization and replacement of a serine at position 65 with a threonine in the chromophore (Chiu et al. 1996).

GFP expression was detected in *P. pruinosa* hypocotyl sections as early as 7 days post infection and persisted throughout the various stages from callus development through rooted plants (Fig. 4 A - C). T0 plants were transferred to soil and grown to maturity in the greenhouse. GFP expression was observed in fruit, husks, seeds, and most notably, embryos exhibiting GFP were in mature seeds (Fig. 4 D - F).

**Figure 4.**
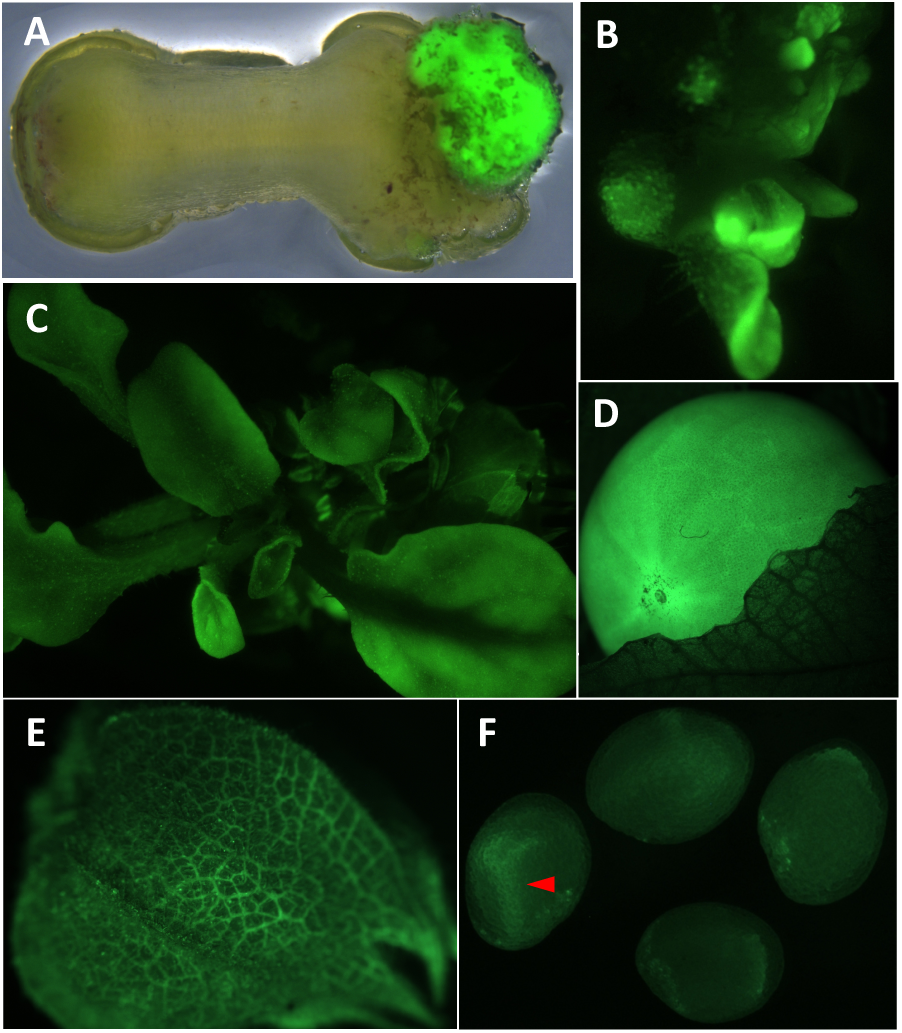
GFP expression in *Physalis pruinosa in vitro* and greenhouse-grown plant material. (*A*) White light and fluorescent overlay image of a hypocotyl with callus 3.5 weeks post infection. (*B*) Shoot regeneration from callus 5 weeks post infection. (*C*) Whole plant. (*D*) Mature fruit. (*E*) Husk. (*F*) Seeds extracted from fruit. Red arrowhead marks embryo in seed.

For further confirmation, PCR analysis with primers designed to detect the *nptII* gene was done on positive control plants recovered from hypocotyl explants that did not undergo infection and GFP-expressing plants recovered from regeneration medium that contained the different kanamycin concentrations. An amplicon of the expected size (735 bp) was detected in all GFP-expressing lines, but not in the controls (Fig. 5).

**Figure 5.**
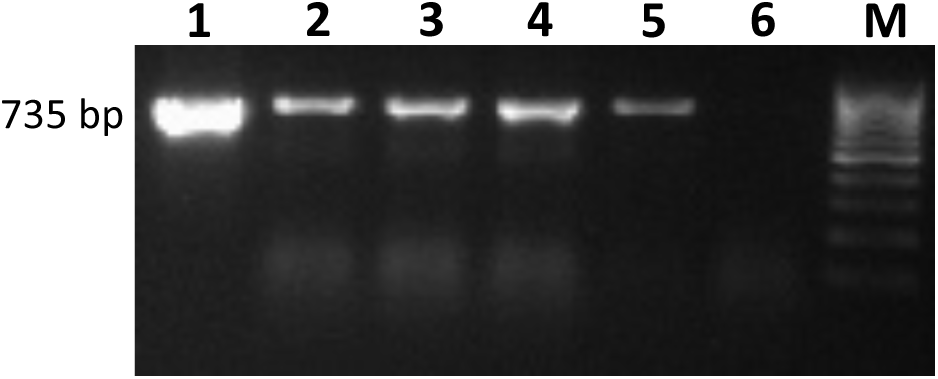
PCR analysis for detection of the *nptII* gene. Expected product size of 735 bp. Lanes: 1: binary plasmid pJL33 DNA, 2 – 5: subset of putative transgenic lines; 6: non-transgenic control, M: molecular size marker (100-bp ladder).

Based on the work reported here, we established a protocol as outlined in Fig. 6 for *Agrobacterium tumefaciens*-mediated transformation of *Physalis pruinosa*. Following infection of hypocotyl explants, it takes approximately 12 weeks for transgenic lines to be at the stage for transfer to soil. All plants transferred to soil survived the process.

**Figure 6.**
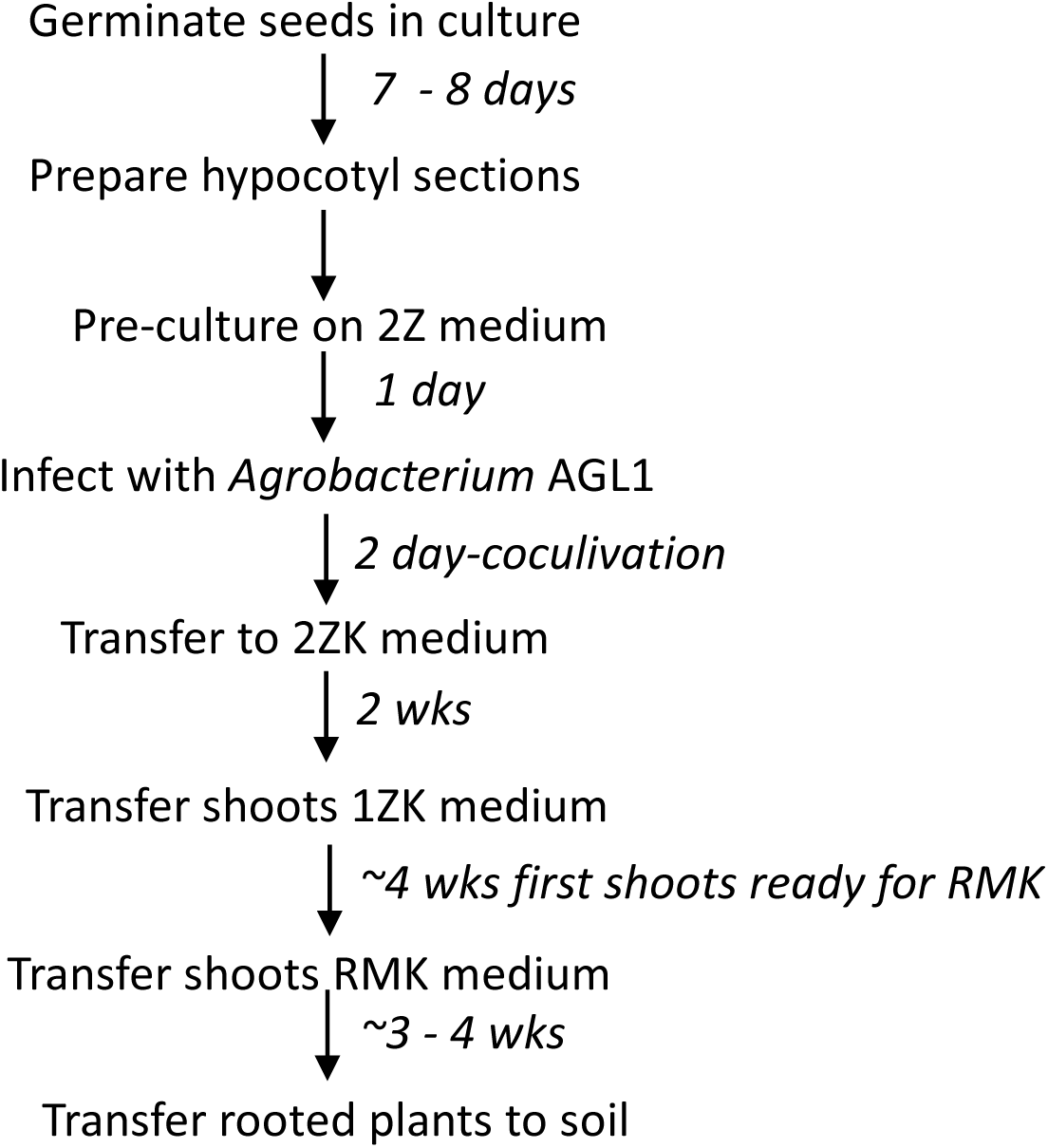
Schematic representation of the optimized *Agrobacterium tumefaciens*-mediated transformation methodology for *Physalis pruinosa*. See Materials and Methods for details on seed sterilization and media composition.

## Discussion

Development of the method reported here was based on our experience with transformation of other Solanaceae family members because although there are reports of plant regeneration and transformation of other *Physalis* species, we were unsuccessful with application of these methods to *P. pruinosa* (Otroshy et al. 2013, Simpson et al. 1995, Van Eck et al. 2018, Wang et al. 2011). As with tomato transformation methodology and reports on transformation of *P. philadelphica*, segments from cotyledons are the explant that works best for *A. tumefaciens*- mediated transformation (Simpson et al. 1995, Van Eck et al. 2018). However, when we investigated plant regeneration from cotyledon and hypocotyl explants we found that only hypocotyl explants regenerated shoots. These findings of the influence of explant choice on plant regeneration support the importance of evaluating different explant sources when developing a gene-delivery method. Hypocotyls from *in vitro*-grown seedlings have also been used for *A. tumefaciens*-mediated transformation of other plant species such as flax and *Brassica napus* (Dong and McHughen 1993, Jonoubi et al. 2005).

Plant selectable marker genes in vectors are key components in development of efficient transformation methods. There are various types of selectable marker genes, but the most commonly used are those that confer resistance to antibiotics and herbicide components, referred to as selection agents, which are incorporated into the culture medium at some determined point following cocultivation. In the past, we performed sensitivity tests to selection agents by incorporation at various concentrations in the culture medium prior to attempting transformation and monitored for response. However, over time we often observed that sensitivity tests performed pre-transformation did not translate into efficient recovery of transgenic lines. Therefore, we changed our approach and now combine preliminary transformation efforts with sensitivity tests to selection agents as reported here for *P. pruinosa*. Through incorporation of kanamycin at five different concentrations in the selective plant regeneration media we identified a concentration that resulted in a 24% transformation efficiency that has been consistent across transformation experiments. The positive effect on transformation efficiency with increasing kanamycin concentration is most likely the result of more efficient inhibition of the growth of non-infected cells that could out-compete infected cells in development. However, as we observed, there is a threshold for the level of kanamycin, in which a high concentration can result in lower transformation efficiency, as evidenced by the decrease in efficiency for *P. pruinosa* hypocotyls cultured on 250 and 300 mg/L. This decrease in efficiency could be explained by *nptII* expression from pJL33 not being high enough to confer resistance at extreme levels. Effect of kanamycin concentration on transformation efficiency has also been reported for other plant species. For rubber tree, (*Heva brasiliensis*) transformation was performed that evaluated media containing different kanamycin concentrations. Results showed that the highest efficiency was obtained on medium containing 250 – 350 mg/L (Jayashree et al. 2003), while concentrations beyond 350 mg/L resulted in a decrease in transformation efficiency. This result was similar to the effect we observed for *P. pruinosa* once the level of kanamycin was increased beyond 200 mg/L.

In addition to selectable marker genes, reporter genes such as GUS and GFP fluorescence proteins allow progress of each step in the development of transformation methods to be monitored. The ability to assess progress at each step helps to identify where effort is needed for efficient recovery of transgenic lines. The reason we chose pJL33 for development of *P. pruinosa* transformation methodology is that, in addition to containing the *nptII* selectable marker gene that we routinely use for other solanaceous species, it also contains the *GFP* reporter gene. One advantage of *GFP* compared to the *GUS* reporter gene is that assessment of GUS expression requires a destructive assay, whereas visualization of GFP expression is done with fluorescent microscopy allowing tracking development throughout different stages. We were able to track progress of recovered transgenic lines, from callus development to plant regeneration through maturity, in the greenhouse. Tracking individual lines from infection of hypocotyls to mature plants would not have been possible with the *GUS* reporter gene.

## Conclusions

To our knowledge, this is the first report of *Agrobacterium tumefaciens*-mediated transformation of *Physalis pruinosa*, also referred to as groundcherry. Transformation methodology was developed for this underutilized *Solanaceae* family member as part of our effort to build resources to improve undesirable agronomic characteristics of *P. pruinosa* to promote its adoption as a specialty fruit crop that has the potential to generate additional income for farmers and provide consumers with a unique fruit crop on a larger scale than would otherwise be possible (Lemmon et al. 2018). Infection of hypocotyl segments from *in vitro*-grown seedlings followed by culture on selective plant regeneration medium containing an optimized concentration of kanamycin is now a routine approach in our lab for recovery of transgenic lines of *P. pruinosa*. Availability of this efficient transformation methodology will greatly facilitate a better understanding of gene function that will contribute to trait improvement in *P. pruinosa*.

## Acknowledgements

The authors acknowledge the National Science Foundation Plant Genome Research Program (IOS-1732253) for support related to their research on *Physalis pruinosa*. The authors thank Mamta Srivastava for her guidance with visualization of green fluorescent protein expression, Daniela Floss for providing pJL33, Esperanza Shenstone for assistance with the manuscript, and Zach Lippman for providing the Rapidase.

